# Identification of lncRNAs associated with early stage breast cancer and their prognostic implications

**DOI:** 10.1101/543397

**Authors:** Arunagiri Kuha Deva Magendhra Rao, Krishna Patel, Sunitha Korivi Jyothi, Balaiah Meenakumari, Shirley Sundersingh, Velusami Sridevi, Thangarajan Rajkumar, Akhilesh Pandey, Aditi Chatterjee, Harsha Gowda, Samson Mani

**Author notes:** Both authors contributed equally to this manuscript (Arunagiri Kuha Deva Magendhra Rao, Krishna Patel). **Correspondence** Samson Mani, Ph. D Associate Professor Department of Molecular Oncology Cancer Institute (WIA), No. 38, Sardar Patel Road, Chennai 600036, India Extn: 131 Harsha Gowda, Ph.D. Faculty Scientist Institute of Bioinformatics 7^th^ floor, Discoverer building, International Tech Park, Bangalore 560 066, India. **E-mail addresses (Complying to the order of author list)**, and.

## Abstract

Breast cancer is a common malignancy among women with the highest incidence rate worldwide. Dysregulation of long non-coding RNAs occurring in the preliminary stages of breast carcinogenesis is poorly understood. In this study, RNA sequencing was done to identify long non-coding RNA expression profiles associated with early-stage breast cancer. RNA sequencing was done in 6 invasive ductal carcinoma (IDC) tissues along with paired normal tissue samples, 7 ductal carcinoma *in situ* (DCIS) tissues and 5 apparently normal breast tissues. We identified 375 differentially expressed lncRNAs (DElncRNAs) in IDC tissues compared to paired normal tissues. Antisense transcripts (∼58%) were the largest subtype among DElncRNAs. About 20% of the 375 DElncRNAs were supported by typical split readings leveraging their detection confidence. Validation was done in n=52 IDC and paired normal tissue by qRT-PCR for the identified targets (ADAMTS9-AS2, EPB41L4A-AS1, WDFY3-AS2, RP11-295M3.4, RP11-161M6.2, RP11-490M8.1, CTB-92J24.3 and FAM83H-AS)1. We evaluated the prognostic significance of DElncRNAs based on TCGA datasets and overexpression of FAM83H-AS1 was associated with patient poor survival. We confirmed that the down-regulation of ADAMTS9-AS2 in breast cancer was due to promoter hypermethylation through in-vitro silencing experiments and pyrosequencing.

## 1. Introduction

Breast cancer is the most common cancer among women (ASR-43.1) with highest mortality rates (Ferlay J). Breast cancer is broadly classified into non-invasive ductal carcinoma *in situ* (DCIS) and invasive-ductal carcinoma (IDC). Understanding the mechanism of breast carcinogenesis at genetic and transcriptional level can aid in characterization of DCIS or early stage IDC tumors. Gene expression signatures are used to classify IDC subtypes of hormone receptor positive (estrogen and progesterone receptors) i.e., luminal A & B and hormone receptor negative-HER2 & basal like (Perou et al., 2000; Sorlie et al., 2001) breast cancer subtypes. Next generation sequencing has enabled global profiling of mRNAs and non-coding RNAs (ncRNAs) including long non-coding RNAs (lncRNAs) and microRNAs. LncRNAs have gained immense importance in gene regulation and are known to play an important role in cancer development and prognosis (Huarte, 2015; Prensner and Chinnaiyan, 2011; Rao et al., 2017). Understanding the divergent expression of lncRNAs in early stage breast tumors can help elucidate its functional role in carcinogenesis.

Specific lncRNA signatures are known to be associated with different molecular subtypes of breast cancer. DSCAM-AS1 was identified specifically in ER positive breast tumors and shown to increase aggression and drug resistance (Niknafs et al., 2016). Similarly, AFAP1-AS1 was predominantly found to be dysregulated in HER2 and triple negative breast cancers (TNBC) (Shen et al., 2015; Yang et al., 2016a). H19 was identified to be over-expressed in ER/PR positive breast adenomas and BC200 was implicated to be distinctly elevated in benign tumors and not in invasive subtypes and hence are of prognostic significance (Adriaenssens et al., 1998; Iacoangeli et al., 2004). HOTAIR was demonstrated to gain activity in BRCA1 mutated tumors. In a normal cell, BRCA1 competes with HOTAIR in binding to EZH2 of PRC2 (Wang et al., 2013). The functional characteristics of certain lncRNAs like UCA1, GAS5 and XIST, have established them as breast cancer associated tumor suppressors while HOTAIR, TINCR and DSCAM-AS1 are known as oncogenic lncRNAs (Wang et al., 2017; Xu et al., 2017). Support vector machine-based prediction of breast cancer intrinsic subtype using lncRNA expression profile and PAM50 gene signature using TCGA datasets was recently proposed as an improved prediction model (Zhang et al., 2018).

Despite known association of lncRNA expression with molecular subtype, recently reported lncRNAs have emerging role in relevant signaling or druggable pathways. LncRNA CYTOR was reported to be associated with breast cancer progression through EGFR signaling pathway (Van Grembergen et al., 2016). NKILA was observed to promote heterotrimeric complex formation (p50/p60/IκB) and inhibit IκB phosphorylation, thus regulating NFκB signaling (Liu et al., 2015). LINK-A was reported to aid in stabilizing HIF1α in normoxic conditions of TNBC. Through BRK/PTK6 activation and phosphorylation of HIF1α, LINK-A substantiates its kinase activation and cancer signaling potential (Lin et al., 2016). Alternatively, breast cancer associated lncRNAs important in drug targeting pathways can also be useful prognostic biomarkers. In the present study, we have done RNA sequencing in early stage tumors (stage I-IIA IDC, DCIS) and non-cancerous breast tissue samples to identify lncRNAs that play a role in early stage breast cancer. We speculate that aberrant expression of lncRNAs could be an early event in breast cancer development and hence the study was aimed to identify dysregulated lncRNAs, and the mechanism of dysregulation in breast cancer.

## 2. Materials and methods

### 2.1 Study population and sample classification

The study cohort includes patients diagnosed and treated for breast cancer at Cancer Institute (WIA), Chennai, Tamil Nadu, India. These patients were histologically confirmed of infiltrating ductal carcinoma (IDC - Stage I-II A) and ductal carcinoma *in situ* (DCIS). Apparently normal breast tissues were obtained from patients undergoing surgery for breast conditions other than malignancy. Samples having >70% for cancer cells following histopathological examination were included in the study. Paired normal and apparently normal tissues completely free of tumor cells were selected and kept frozen (−80°C) until further processing. Total RNA sequencing was done for 24 samples i.e. tumor (n=6), paired normal (matched normal; n=6), DCIS (n=7), and apparently normal (n=5). Validation cohort of IDC (n=52) and corresponding paired normal tissue were used to gauge candidate lncRNAs. The clinico-pathological features of patients in the discovery and validation cohort are detailed in Supplementary Table S1. All patients were informed about the study and their written consent for participation was obtained. The Institutional Ethical Committee approved the study and the protocol.

### 2.2 RNA isolation and library preparation

Total RNA was isolated from frozen tissues using TRIZOL method and purification by Nucleospin RNA isolation kit (Machery-Nagel, GmbH), which includes an on-column DNase treatment. The quality and quantity of total RNA was evaluated through Bioanalyzer 2100 (Agilent Technologies, CA, USA). Ribosomal RNA was depleted (Epigentek, USA) and cDNA library was prepared using Illumina TruSeq Stranded Total RNA Library Prep Kit. The library profile was verified using 2100 Bioanalyzer (Agilent Technologies, CA, USA). Subsequent RNA sequencing of cDNA libraries with paired-end reads (2 x 100 bps reads) were performed according to the standard Illumina protocol using HiSeq2500 sequencing platform.

### 2.3 RNA sequencing and data analysis

Raw reads were assessed for Phred quality using FastQC (Andrews); and low bases and adaptor sequences were trimmed off using Fqtrim (Pertea, 2015) retaining reads of length ≥ 75 bases. Clean reads were aligned against human reference genome (GRCh38 assembly) with Gencode V24 annotation using Hisat2 (Baruzzo et al., 2017) with default parameters. Exon centric read counts were obtained from binary alignment map (BAM) using HTSeq(Anders et al., 2015) using the script ‘htseq count’ for all samples independently. LncRNAs identified with ≥ 15 reads in at least 3 samples per cohort i.e. IDC, paired normal, DCIS and apparent normal were further investigated for differential expression using DESeq (Anders and Huber, 2010). Read counts obtained from HTSeq were normalized using ‘estimateSizeFactors’ variance and were modeled using ‘estimateDispersions’. The differentially expressed genes were computed using ‘nbinomTest’ functions of DEseq. Significant differential expression was defined if |log_2_ (fold-change)| > 1 and q-value (BH adjusted P value) < 0.1. Expression profile of long non-coding RNA from TCGA breast cancer dataset (TCGA-BRCA; n=837 invasive tumors and n=105 normal samples) was used for survival analysis (Li et al., 2015). Kaplan-Meier plots for differentially expressed lncRNAs were generated for tumor stages as well as molecular subtypes and evaluated using log rank test.

### 2.4 LncRNA-mRNA co-expression network analysis

Pearson’s correlation coefficient (PCC) was used to determine linear correlation between mRNA and lncRNA expression profiles using R. Differentially expressed lncRNA-mRNA pairs with |PCC| ≥ 0.9 were considered for network analysis using STRING v10 (Szklarczyk et al., 2015) with organism “Human” as backend database and Cytoscape (Shannon et al., 2003).

### 2.5 Real-time quantitative PCR

Total RNA of 500 ng was used for preparing cDNA libraries using QuantiTect Reverse Transcription Kit (Qiagen, USA). Gene expression was estimated by QuantStudio 12K Flex Real-Time PCR System (Applied Biosystems, USA) using TaqMan™ gene expression assays (Applied Biosystems, USA) containing primers and probes specific for lncRNA and GAPDH. The expression values were calculated using the 2^-ΔCt^ method (ΔCt = ΔCt target gene-ΔCt reference gene).

### 2.6 siRNA mediated knock-down of DNMT1

Expression of ADAMTS9-AS2 was evaluated in MDAMB-231 and MCF7 cells. The cells were cultured in DMEM with 10% fetal bovine serum at 37°C. Knockdown was carried out using Lipofectamine 3000 (Invitrogen, USA), siRNA targeting DNMT1 (Ambion, USA) with cells maintained in OptiMEM (Life Technologies, USA) during and after transfection. Transfected cells were collected after 48 hours and 72 hours for total RNA and DNA isolation.

### 2.7 DNA extraction, Bisulfite treatment and pyrosequencing

Genomic DNA was extracted from tissues and cultured MDAMB-231 and MCF7 cells using Nucleospin Kit (Machery and Nagel, GmbH). About 500 ng of DNA was used for bisulfite treatment following manufacturer’s protocol of EZ DNA Methylation-Gold Kit (Zymo Research, CA, USA). Bisulfite treated DNA was amplified using inventoried PyromarkCpG assay*Hs_AC132007.1_01_PM* (Qiagen, GmbH) with primers spanning ADAMTS9-AS2 promoter region. The amplified fragment was sequenced using Pyromark Q48 Autoprep (Qiagen, GmbH) and analyzed by PyroMark Q24 Software v 2.0.7.

### 2.8 Statistical analyses

GraphPad Prism (Version 7.0, La Jolla, California, USA) was used for evaluating qRT-PCR gene expression data. Student’s t-test was used for pair-wise analysis of tumor and paired normal samples. Welch correction was done if significant difference in variance was observed and Wilcoxon rank sum test was applied whenever non-Gaussian distribution was followed.

## 3. Results

### 3.1 Expression profile of lncRNAs in ductal carcinoma in situ and invasive ductal carcinoma

RNA sequencing resulted in generation of ∼89 million reads per sample with ∼87.24% alignment against human genome build Hg38. We identified ∼2,689 lncRNAs and ∼18,132 mRNAs with ≥ 15 reads in at least 3 samples per cohort [Table 1, Supplementary Table S 2]. In agreement with previous reports, lncRNAs were expressed at comparatively lower levels than mRNAs [Supplementary Figure 1A-D]. Principal component analysis (PCA) plots based on lncRNA quantification showed distinct segregation of tumors (IDC and DCIS) from paired and apparent normal samples reflecting the characteristic variation of lncRNA expression profile [Figure 1A, Supplementary Figure 1E]. Differential expression analysis was performed between IDC, DCIS and control samples in four categories i.e., IDC *vs* paired normal (TN), IDC *vs* apparent normal (TA), DCIS *vs* apparent normal (DA) and IDC *vs* DCIS (TD)[Figure 1B-D].

**Table 1.** Number of differentially expressed lncRNAs in ductal carcinoma *in-situ* and early stage breast cancer

| Comparison set | lncRNA |  |  |  |
| --- | --- | --- | --- | --- |
|  | Overexpressed | Down regulated | Total | Split reads |
| **IDCvs. Paired normal** | 195 | 180 | 375 | 96 |
| **IDCvs. Apparent normal** | 38 | 56 | 94 | 25 |
| **DCIS vs. Apparent normal** | 29 | 40 | 69 | 24 |
| **IDCvs. DCIS** | 5 | 7 | 12 | 3 |

**Figure 1.**
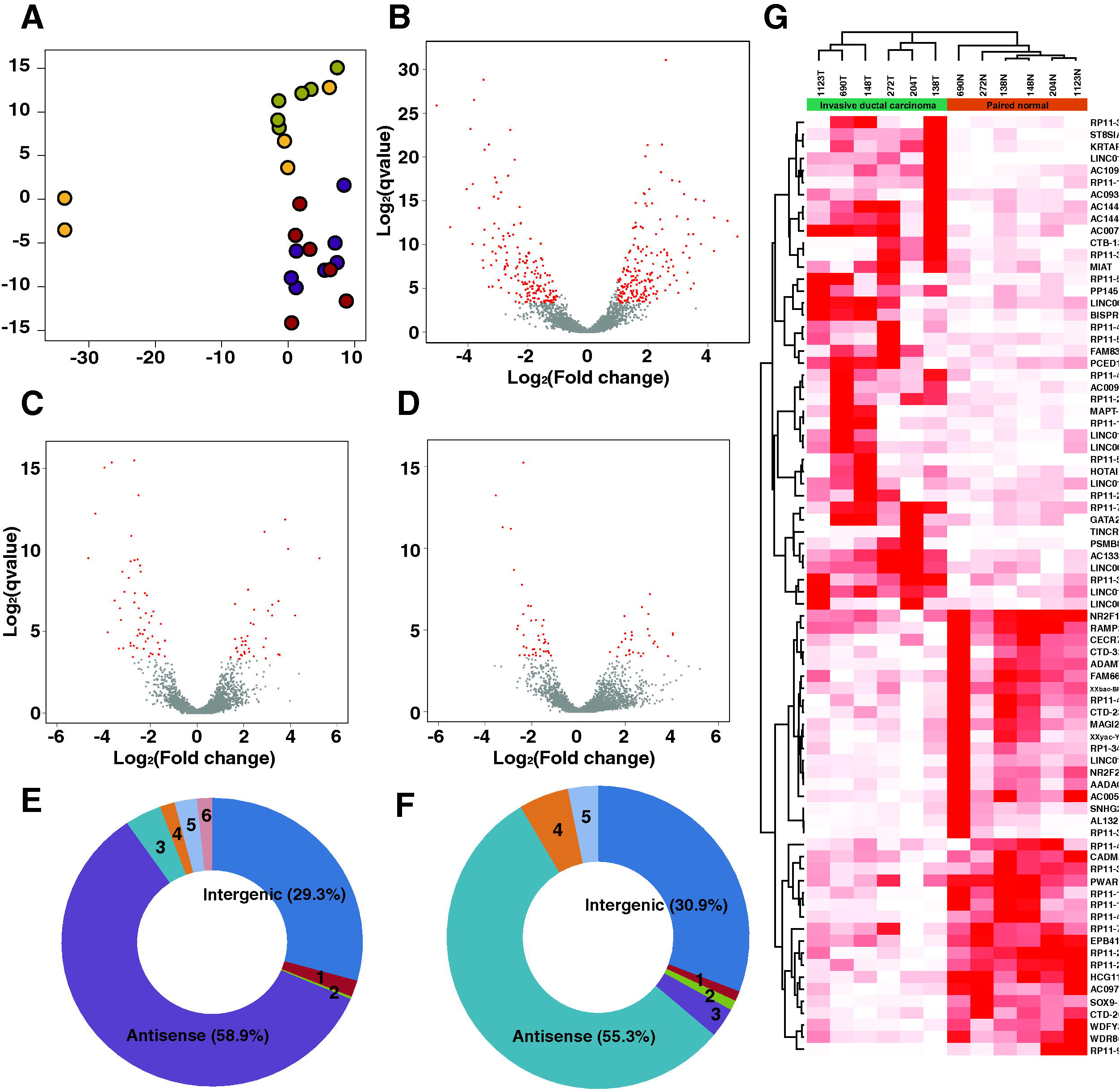
Summary of differentially expressed lncRNAs identified in ductal carcinoma *in-situ* and early stage breast cancer. **(A)**. Principal component analysis based on lncRNA expression profileto demonstrate distinct segregation of tissues of various pathological types. Color legend. Apparent normal: Yellow, DCIS: Purple samples, Paired normal: Green, IDC: Red**(B)**Volcano plot represents the expression pattern of lncRNA in IDC*vs.* paired normal samples. **(C)**Volcano plot represents the expression pattern of lncRNA in IDC*vs.* apparent normal samples. **(D)**Volcano plot represents the expression pattern on lncRNA inDCIS*vs.* apparent normal samples. **(E)** Pie chart representing DElncRNA subtypes inIDC *vs.* paired normal samples[1-Intron overlapping (1.9%); 2-Non-coding transcript (0.3%); 3-TEC (0.4%); 4-Sense overlapping (1.5%); 5-Processed transcript (2.4 %); 6-Completely intronic (1.6 %)].**(F)**Pie chart representing DElncRNA subtypes inIDC *vs.* apparent normal samples [1-Intron overlapping (1.1 %); 2-Completely intronic (1.1 %); 3-TEC (3.2 %); 4-Processed transcript (5.3 %); 5-Sense overlapping (3.2 %)].**(G)**Heatmap with supervised clustering represents the expression trend of DElncRNAs in IDC*vs*. paired normal samples. **(H)**Heatmap with supervised clustering represents the expression trend of DElncRNAs in IDC*vs*. apparent normal samples. **(I)**Heatmap with supervised clustering represents the expression trend of DElncRNAs in DCIS *vs*. apparent normal samples.

We observed antisense RNAs (asRNA) and long intergenic RNAs (lincRNAs) to be the major lncRNA subtypes differentially expressed among these four groups. Antisense RNAs accounted for 58.9% of total differentially expressed lncRNAs in IDC compared to paired normal and 55.3% compared to apparently normal samples. [Figure 1 E-F]. WDR86-AS1, emerged as a novel antisense lncRNA in our data whereas ADAMTS9-AS2 (Li et al., 2017; Peng et al., 2017) and ST8SIA6-AS1 (Yang et al., 2016a; Yang et al., 2016b) have previously been reported in other studies [Figure 1 G-H].

### 3.2 Identification of novel lncRNAs differentially expressed in breast tumors

Dysregulated lncRNAs with evidence of ≥ 2 junction reads in each comparison groups were further investigated [Supplementary Figure 1F-I]. We identified 21 lncRNAs (eleven overexpressed and ten down regulated) showing a differential expression pattern [Table 2, Figure 2]. Among them, MIAT, FAM83H-AS1, EPB41L4A-AS1, WDFY3-AS2 and RP11-392O17.1 were commonly deregulated in TN, TA and DA comparison groups [Figure 2]. Further, LINC01614, RP11-490M8.1 and CTB-92J24.3 were novel DElncRNAs identified in early staged breast cancer.

**Table 2.** List of differentially expressed lncRNAscommon among various comparison sets

| lncRNA | IDC vs.<br>Apparent<br>normal | IDC vs.<br>Paired<br>normal | DCIS vs.<br>Apparent<br>normal | Expression status |
| --- | --- | --- | --- | --- |
| MIAT | 2.89 | 1.47 | 2.72 | Overexpressed |
| FAM83H-AS1 | 1.96 | 1.92 | 2.01 | Overexpressed |
| LINC01614 | 5.24 | 6.1 | - | Overexpressed |
| RP11-527N22.1 | 4.2 | 3.77 | - | Overexpressed |
| TINCR | 3.22 | 4.22 | - | Overexpressed |
| CTB-131K11.1 | 2.42 | 1.96 | - | Overexpressed |
| RP11-126H7.4 | 2.22 | 1.77 | - | Overexpressed |
| LINC01105 | 3.48 | 4.04 | - | Overexpressed |
| AC093642.3 | 2.94 | 3.39 | - | Overexpressed |
| ST8SIA6-AS1 | - | 2.48 | 3.21 | Overexpressed |
| AC109826.1 | - | 2.12 | 2.99 | Overexpressed |
| RAMP-AS1 | -1.38 | -1.43 | - | Downregulated |
| ADAMTS9-AS2 | -1.65 | -3.31 | - | Downregulated |
| RP11-490M8.1 | -2.32 | -1.8 | - | Downregulated |
| RP11-92A5.2 | -3.53 | -5.05 | - | Downregulated |
| EPB41L4A-AS1 | -1.55 | -1.18 | -1.5 | Downregulated |
| WDFY3-AS2 | -1.68 | -1.44 | -1.65 | Downregulated |
| RP11-392O17.1 | -2.69 | -2.72 | -2.63 | Downregulated |
| RP11-161M6.2 | -2.44 | -2.11 | - | Downregulated |
| CTB-92J24.3 | -2.42 | -2.42 | - | Downregulated |
| RP11-295M3.4 | - | -2.79 | -2.77 | Downregulated |

**Figure 2.**
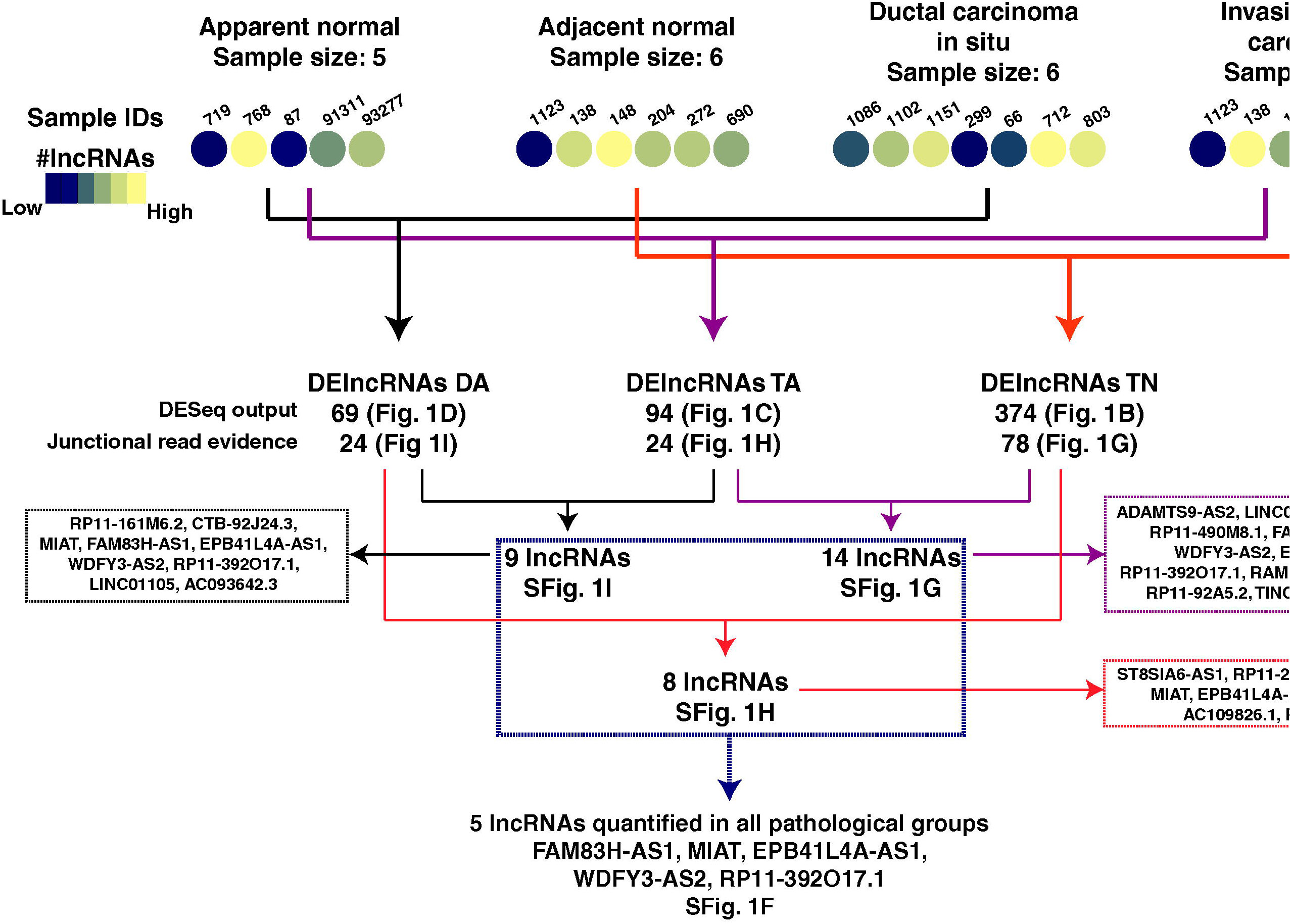
Schematic of lncRNA analysis and cross-comparison of differentially expressed lncRNAs in multiple comparison groups.

### 3.3 Validation of candidate lncRNA expression in breast tumor and paired normal

We selected 12 candidate lncRNAs (5 Upregulated lncRNAs: MIAT, FAM83H-AS1, LINC01614, ST8SIA6-AS1, CTB-131K11.1 and 7 down regulated lncRNAs: ADAMTS9-AS2, EPB41L4A-AS1, WDFY3-AS2, RP11-161M6.2, RP11-295M3.4, RP11-490M8.1, CTB-92J24.3) for validation using TaqMan™ gene expression assays in n=52 early staged IDC samples [Figure 3A]. We observed statistically significant dysregulation of seven out of twelve lncRNAs identified using RNA-Seq. Among them ADAMTS9-AS2 [Figure 3B] was observed to be the most commonly down regulated lncRNA in tumor tissues (13.59 folds). We also confirmed significant down regulation of CTB92J24.3(11.82 folds), RP11-295M3.4 (3.5 folds), RP11-490M8.1 (3.7 folds), WDFY3-AS2 (4.3 folds) and EPB41L4A-AS1 (2.09 folds) [Figure 3C-G]. FAM83H-AS1 was most significantly overexpressed lncRNA in tumors (8.9 folds) compared to the paired normal tissues [Figure 3H]. Although, MIAT and LINC01614 were upregulated, statistically were insignificant [Figure 3I-J]. Whereas, ST8SIA6-AS1 and CTB-131K11.1 were found to be down regulated contradicting out RNA sequencing results [Figure 3K-L]. To evaluate the involvement of receptor status, expression levels of 12 DElncRNAs from validation cohort were correlated with receptors status (ER, PR, HER2) [Supplementary Figure 4A-D]. We observed that MIAT was overexpressed exclusively in samples that were ER + PR + Her2^+^ whereas RP11-161 M6.2 was overexpressed in ER^-^PR^-^.

**Figure 3.**
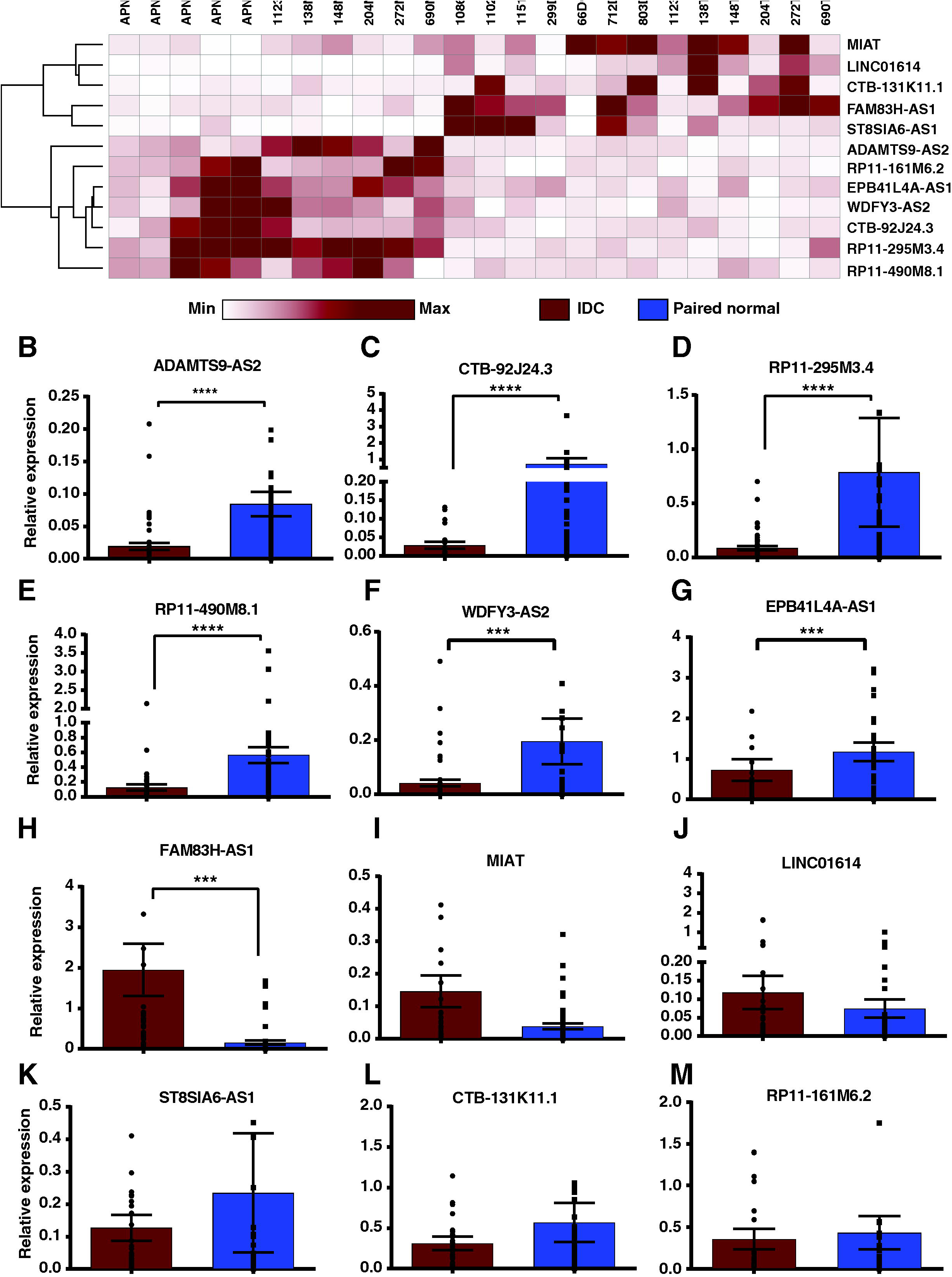
Expression validation of differentially expressed lncRNAs using qRT-PCR in cohort of 52 early stage breast cancer samples. **(A)** Heatmap of differentially regulated showing expression trend in discovery set of samples. **(B)**Relative expression of ADAMTS9-AS2 **(C)**Relative expression of CTB-92J24.3 **(D)**Relative expression of RP11-295M3.4 **(E)**Relative expression of RP11-490M8.1 **(F)**Relative expression of WDFY3-AS2 **(G)**Relative expression of EPB41L4A-AS1 **(H)**Relative expression of FAM83H-AS1 **(I)**Relative expression of MIAT **(J)**Relative expression of LINC01614 **(K)**Relative expression of ST8SIA6-AS1 **(L)**Relative expression of CTB-131K11.1 **(M)**Relative expression of RP11-161M6.2 [B-M are relative expression levels of lncRNA evaluated in validation set of samples]; (Wilcoxon sign rank test p-value < 0.0001= ****, p<0.001= *** and not indicated for non-significant candidates).

**Figure 4.**
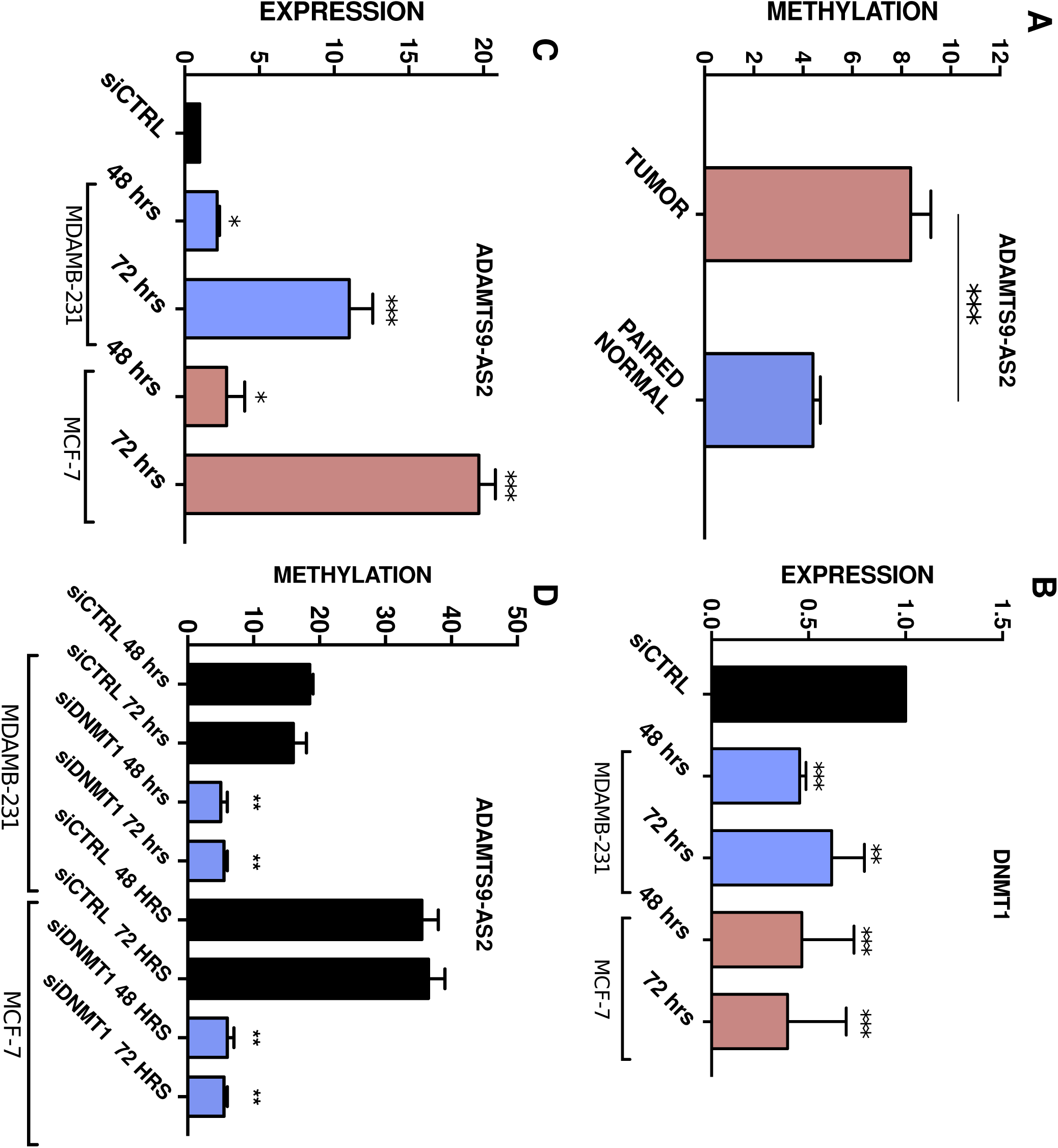
**(A)**Relative methylation levels of ADAMTS9-AS2 promoter in tumor vs paired normal tissue [N=52] **(B)**Expression levels of DNMT1 with siRNA treatment in MDAMB231and MCF7 cells. **(C)** Expression of ADAMTS9-AS2 in MDAMB-231 and MCF7 cells onDNMT1 knock-down **(D)** Relative methylation levels of ADAMTS9-AS2 promoter in MDAMB-231 and MCF7 cells with DNMT1 knock-down [***=p<0.001, ** = p<0.01 & *=p<0.05.

### 3.4 ADAMTS9-AS2 promoter is hyper-methylated in breast tumors

Yao *et al* reported the downregulation of ADAMTS9-AS2 by promoter methylation in gliomas (Yao et al., 2014). Hence methylation levels of the promoter region of ADAMTS9-AS2 in our validation set of tumor and paired normal samples (n= 52) was done using pyrosequencing. We observed a nearly two folds (1.9) increase in methylation levels (p< 0.0001) in the promoter region (+879 to +929 bp from TSS) of tumor samples compared to paired normal samples [Figure 4A].

### 3.5 Knock-down of DNA methyltransferase 1 increases ADAMTS9-AS2 expression

In order to investigate promoter methylation mediated regulation of ADAMTS9-AS2 expression, DNMT1 was knocked down in MDAMB-231 and MCF7 using short interfering RNA. The down regulation of DNMT1 led to subsequent over expression of ADAMTS9-AS2 by 1.93-fold (p<0.001) and 2.32-fold (p<0.001) in MDAMB-231 and MCF7 respectively [Figure 4B-C]. Loss of promoter methylation was observed using pyrosequencing in DNMT1 siRNA transfected MDAMB-231 (2.6 folds; p=0.001) and MCF-7 cells (6.7 folds; p=0.007) [Figure 4 D]. These results show that ADAMTS9-AS2 is over expressed in both MDAMB-231 and MCF7 cells following DNMT1 silencing indicating methylation-mediated suppression of ADAMTS9-AS2 in breast cancer cells.

### 3.6 Prognostic lncRNAs in early stage breast cancer

Survival analysis was done to investigate the prognostic potential of candidate lncRNA using TCGA datasets. We observed FAM83H-AS1 was significantly overexpressed by ∼4 fold in TN, TA as well as DA pairs and its overexpression is associated with overall poor survival in luminal A, ER positive tumors, stage 3 datasets and overall breast tumor datasets irrespective of subtypes [Figure 5A-D]. Overexpression of WDFY3-AS2 in luminal A, ER positive tumors and breast tumor datasets irrespective of subtypes [Figure 5 E, F and H] is significantly associated with adverse outcomes. Whereas, down regulation of RP11-161M6.2 in breast cancer and CTB-92J24.3 in stage 3 was observed significantly associated with poor overall survival [Figure 5K]. We observed significant association with overexpression of WDFY3-AS2 [Figure 5G] and down regulation of RP11-161M6.2 in stage2 of breast cancer based on TANRIC analysis indicating them as potential early prognostic markers [Figure 5G and J].

**Figure 5.**
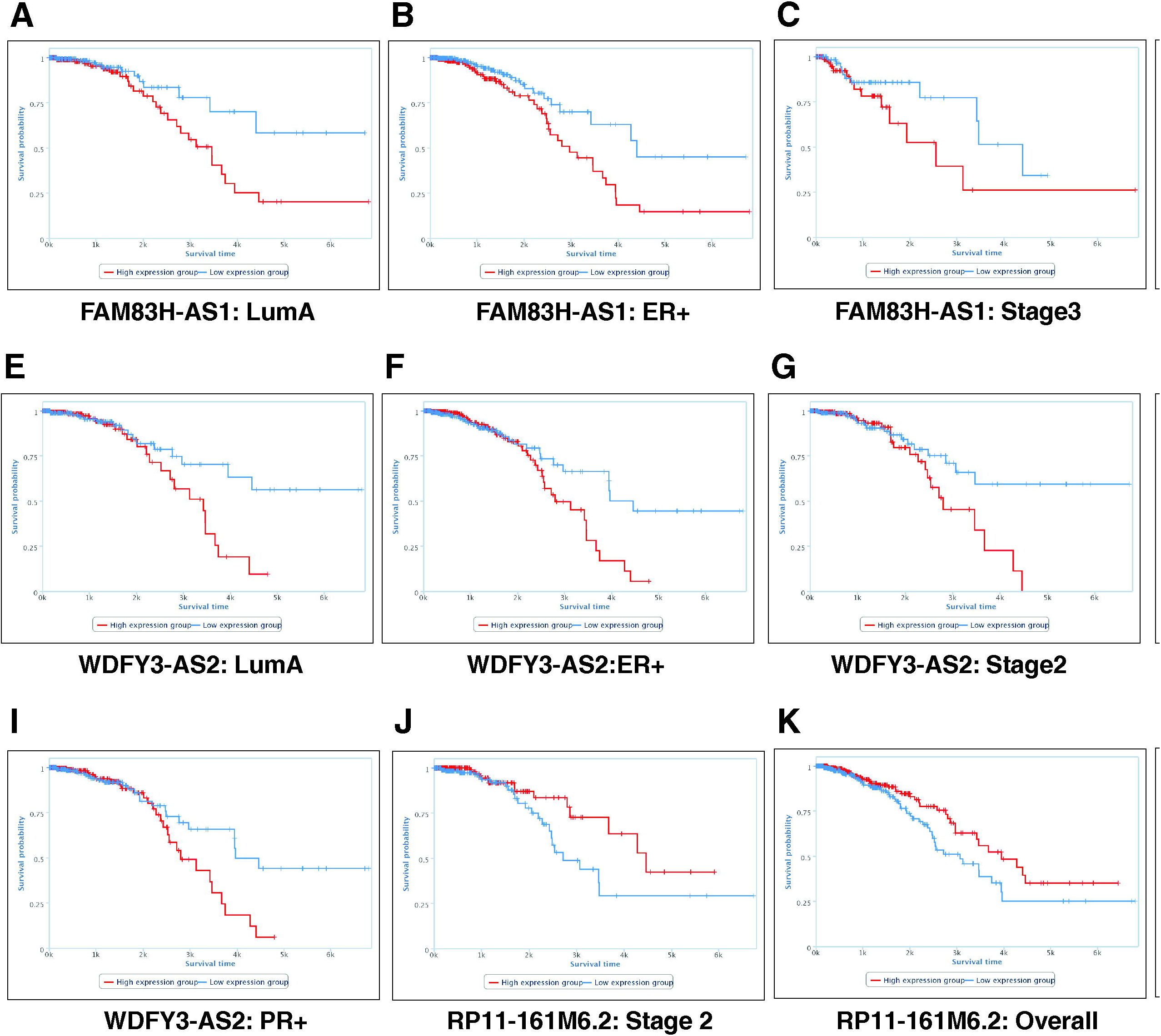
Kaplan Meir plots derived from TANRIC depicting significant overall poor survival of patients associated with differentially expressed lncRNAs. **(A)** FAM83H-AS1 in Luminal A molecular subtype **(B)** FAM83H-AS1 in ER+ molecular subtype **(C)** FAM83H-AS1 in Stage 3 dataset **(D)** FAM83H-AS1 in overall breast cancer dataset **(E)** WDFY3-AS2 in Luminal A molecular subtype **(F)** WDFY3-AS2 in ER+ molecular subtype **(G)** WDFY3-AS2 in Stage 2 dataset **(H)** WDFY3-AS2 in overall breast cancer dataset **(I)** WDFY3-AS2 in PR+ molecular subtype **(J)** RP11-161M6.2 in stage 2 dataset **(K)** RP11-161M6.2 in overall breast cancer dataset **(L)** CTB-92J24.3 in stage 3 dataset.

### 3.7 Co-expression and pathway analysis

Guilt-by-association method was employed to speculate the putative functions of lncRNAs. This approach investigates the association of mRNA expression patterns with lncRNAs using Pearson correlation analysis. A correlation analysis between DElncRNA-DEmRNA pairs was done and only those with Pearson correlation coefficient (PCC) ≥ |0.9| were considered significantly co-expressed. The co-expressed pairs were filtered for lncRNA with typical junctional read evidence which led to the identification of 2,398 pairs consisting of 78 lncRNA and 1,097 mRNA between IDC and paired normal samples and 385 pairs consisting of 24 lncRNA and 245 mRNA between IDC and apparent normal samples.

Similarly, 26 pairs were co-expressed in DCIS *vs.* apparent normal samples consisting of 11 lncRNA and 26 mRNA and 10 co-expressed lncRNA-mRNA pairs in IDC compared to DCIS representing 3 lncRNA and 10 mRNA [Supplementary Table S7-10]. Among, 2,398 co-expressed lncRNA-mRNA pairs in IDC *vs* paired normal samples, 2,225 (92.83%) harbors on different chromosomes (trans-acting) whereas remaining pairs are cis-acting. Similarly, 351 (91.64%) out of 383 in IDC *vs* apparent normal samples and 23 (85.17%) out of 27 in DCIS *vs.* apparent normal samples are located on different chromosomes.

Co-expressed mRNAs were further analyzed using StringDB for network analysis. To augment guilt by association concept, we further focused on mRNA network that are reported to co-express irrespective of lncRNA. We observe that partial sets of mRNAs from 22 DElnRNAs in IDC compared to paired normal samples were co-expressed according to StringDB analysis. After removing disconnected nodes and filtering high confidence nodes from the network, genes co-expressed with RP11-142C4.6 [Supplementary Figure 5A] were found enriched for extracellular regions (red nodes) and overrepresented for extracellular matrix organization (green nodes) and disassembly (blue nodes) whereas genes co-expressed with RAMP2-AS1were enriched on the cell membrane (red nodes) [Supplementary Figure 5A and B]. Genes co-expressed with RP11-701H24.4 were enriched for integral component of membrane (green nodes) and activation of cellular processes (blue nodes) [Supplementary Figure 5C]. In case of PSMB8-AS1, we observed overrepresentation of immune response and (red nodes) involved in type I interferon-signaling pathway (blue nodes) [Supplementary Figure 5D]. We observed enrichment of biological process like, cell division (yellow nodes), cell cycle process (pink nodes) and microtubule cytoskeleton (red nodes) in genes positively co-expressed with TINCR and negatively co-expressed with LINC01359 [Supplementary Figure 6 and 7]. Interestingly, most genes co-expressed with PSMB8-AS1, TINCR and LINC01359 are also known to co-express with each other according to StringDB. Using Cytoscape, we were able to segregate the sub network of 76 genes potentially governed jointly by TINCR (65 genes) and LINC01359 (55 genes), which resulted in sub modules of genes with core histone protein domains (green nodes) and involved in pathways in cancer (blue nodes).

## 4. Discussion

Aberrant expression of long non-coding RNAs (lncRNAs) is documented in various cancers (Huarte, 2015; Prensner and Chinnaiyan, 2011). In recent years, lncRNAs have gained importance in early detection and better prognosis of tumors (Chandra Gupta and Nandan Tripathi, 2017). Although several lncRNAs associated with breast cancer have been reported previously, studying aberrantly expressed lncRNAs specific to early stage breast cancer will provide insight into molecular mechanisms associated with breast cancer development. It will also result in identification of putative markers that might be useful in diagnosis or prognosis of breast cancer. Previous studies have associated altered expression of lncRNAs with specific breast cancer subtypes. For example, HOTAIR is a lncRNA that is highly expressed in HER2+ breast cancers whereas HOTAIRM1 is highly expressed in basal-like subgroup of breast cancers (Su et al., 2014). LuminalA types showed over expression of LINC00160 and abundance of DSCAM-AS1 was reported in luminalB subtypes of breast cancer (Jonsson et al., 2015; Vu et al., 2016). MALAT, lncRNA-ATB, BC200, XIST, H19 are some of other lncRNAs frequently associated with breast tumorigenesis and progression (Hansji et al., 2014). Functionally important lncRNAs in early stage breast cancers are less reported. Our study evaluated the landscape of lncRNA expression in early stage breast cancer [IDC (Stage I-IIA) and DCIS breast tissues] to identify aberrantly expressed lncRNAs.

The DESeq analysis resulted in identification of 375 DElncRNAs in IDC compared to paired normal samples and 94 DElncRNAs in IDC compared to apparent normal samples. The analysis also identified 69 DElncRNAs in DCIS compared to apparent normal samples. We identified several antisense lncRNAs including ADAMTS9-AS2, EPB41L4A-AS1, WDFY3-AS2, FAM83H-AS1, ST8SIA6-AS1, CTB-92J24.3 and CTB-131K11.1 that were aberrantly expressed. Twelve candidate lncRNAs that showed significant differential expression were further validated in 52 paired tumor and normal breast samples. We observed significant down regulation of ADAMTS9-AS2, WDFY3-AS2, RP11-295M3.4, RP11-490M8.1, CTB-92J24.3 and significant over expression of FAM83H-AS1 in breast cancer. We found ADAMTS9-AS2 to be significantly down regulated in tumor compared to paired normal breast tissues. ADAMTS9-AS2, is an antisense transcript originating from the opposite stand coding for ADAMTS9 which is a known inhibitor of angiogenesis and is implicated to have a tumor suppressive role. Functional importance of ADAMTS9 in nasopharyngeal and esophageal cancers has been described (Lo et al., 2010). ADAMTS9-AS2 like ADAMTS9 is down regulated in glioblastoma (Yao et al., 2014), colorectal cancer(Li et al., 2016), bladder cancer, lung adenocarcinoma and ER+ breast cancers(Li et al., 2017). Yao *et al* have shown that promoter methylation regulatesADAMTS9-AS2 expression by knocking down DNMT1 in glioma cells. We found that methylation of ADAMTS9-AS2 controls its expression through correlative DNMT1 knock-down in MDAMB231 and MCF7 cells. Similar results were observed when methylation levels at ADAMTS9-AS2 promoter were compared between tumors and paired normal tissues using pyrosequencing. We observed DNA methylation mediated loss of ADAMTS9-AS expression in stage I breast cancer. Among other down regulated lncRNAs, WDFY3-AS2 has recently been reported with TGF-B induced EMT of breast cancer cells through hnRNP-R modulated positive regulation of STAT3 and WDFY3 (Richards et al., 2016). Down regulation of WDFY-AS2 was found in diffuse glioma and strongly associated with poor prognosis (Wu et al., 2018). EPB41L4A-AS1 (also known as TIGA1) has been shown to be transcribed during growth arrest but has not been extensively studied in cancer to elucidate its role (Yabuta et al., 2006). RP11-161 M6.2 was found to be over expressed in ER/PR negative and HER2 positive breast cancers in our samples. The finding indicates an association of RP11-161 M6.2 and estrogen receptor and is possibly down regulated in estrogen mediated signaling. Similarly, MIAT was dominantly expressed in ER/PR/HER2+ breast cancers samples.

FAM83H-AS1 was consistently over expressed in breast tumor samples and overall survival analysis of TCGA data sets showed poor prognosis of the up regulated group which are in agreement with other studies in breast, colorectal and lung cancer (Lu et al., 2018; Yang et al., 2016a; Yang et al., 2016c; Zhang et al., 2017). Functional studies have demonstrated that knock-down of FAM83H-AS1 proliferative potential through MET/EGFR signaling in lung adenocarcinoma and NOTCH1 signaling pathway in colorectal cancer. Overexpression of FAM83H-AS1 in luminal type breast cancer associated with good prognosis in patients (Yang et al., 2016a). Detection of FAM83H-AS1 expression levels in plasma could be a potential diagnostic and prognostic biomarker for breast cancer.

In summary, this study has shed light on novel lncRNA and substantiated several previous findings on lncRNA involved in early stage breast cancers. We report 375 and 94 lncRNA differentially expressed in tumor samples compared to paired and apparent normal samples respectively and 69 DElncRNAs in DCIS compared to apparent normal samples. Seven down regulated and five upregulated lncRNA were further validated to discover significant lncRNA candidate with potential role in breast carcinogenesis. ADAMTS9-AS2 was one of the lncRNA consistently down regulated in patient samples and experimental evidence proved promoter methylation as major cause of ADAMTS9-AS2 down regulation in breast cancer. Moreover, LINC01614, RP11-490M8.1 and CTB-92J24.3are novel lncRNA reported in our study that has not been associated with breast cancer earlier. Our study also contributes to the existing evidence on MIAT and FAM83H-AS1 as crucial lncRNA expressed at preliminary stages of breast cancer

## Data availability

Raw sequencing data is available in Sequence Read Archive hosted by National Center for Biotechnology Information Search database (NCBI) with accession number PRJNA484546.

## Acknowledgment

We thank Dr. Uma Devi K.R. and Dr. S. Sivakumar, National institute for Research in Tuberculosis for providing pyrosequencing facility. Krishna Patel is recipient of Senior Research Fellowship from Council of Scientific and Industrial Research (CSIR).

## Supplementary tables legends

**Supplementary Table S1.** List of clinicopathological features of patients’ tissue samples used in discovery and validation cohort in the study.

**Supplementary Table S2.** Read alignment statistics and number of genes identified in different samples.

**Supplementary Table S3.** Complete list of differentially expressed lncRNAs identified to be differentially expressed in IDC (T) *vs.* paired normal (N) samples with adjusted p-values <0.1 in this study along with normalized read counts from individual samples.

**Supplementary Table S4.** Complete list of differentially expressed lncRNAs identified to be differentially expressed in IDC (T) *vs.* apparent normal (APN) with adjusted p-values <0.1 in this study along with normalized read counts from individual samples.

**Supplementary Table S5.** Complete list of differentially expressed lncRNAs identified to be differentially expressed in DCIS *vs.* apparent normal (APN) with adjusted p-values <0.1 in this study along with normalized read counts from individual samples.

**Supplementary Table S6.** Complete list of differentially expressed lncRNAs identified to be differentially expressed in IDC (T) *vs.* DCIS with adjusted p-values <0.1 in this study along with normalized read counts from individual samples.

**Supplementary Table S7.** Complete list of dysregulated mRNA co-expressed with dysregulated lncRNAs supported by split reads in IDC *vs.* paired normal with Pearson correlation coefficient (PCC) ≥ 0.9.

**Supplementary Table S8.** Complete list of dysregulated mRNA co-expressed with dysregulated lncRNAs supported by split reads in IDC *vs.* apparent normal with Pearson correlation coefficient (PCC) ≥ 0.9.

**Supplementary Table S9.** Complete list of dysregulated mRNA co-expressed with dysregulated lncRNAs supported by split reads in DCIS *vs.* apparent normal with Pearson correlation coefficient (PCC) ≥ 0.9.

**Supplementary Table S10.** Complete list of dysregulated mRNA co-expressed with dysregulated lncRNAs supported by split reads in IDC *vs.* DCIS with Pearson correlation coefficient (PCC) ≥ 0.9.

**Supplementary Table S 11.** List of gene expression assays

## Supplementary figure legends

**Supplementary Figure 1. Expression pattern of lncRNAs and protein coding genes in various pathological subtype and comparison of DElncRNAs in different groups.(A)** Comparative histogram represents relatively lower expression of lncRNAs (blue bars) compared to protein coding genes (grey bars) based on raw read count profile of apparent normal samples (n=5) **(B)** Comparative histogram represents relatively lower expression of lncRNAs (green bars) compared to protein coding genes (grey bars) based on raw read count profile of paired normal samples (n=6) **(C)** Comparative histogram represents relatively lower expression of lncRNAs (yellow bars) compared to protein coding genes (grey bars) based on raw read count profile of DCIS samples (n=7) **(D)** Comparative histogram represents relatively lower expression of lncRNAs (red bars) compared to protein coding genes (grey bars) based on raw read count profile of IDC samples (n=6) **(E)** Principal component analysis using normalized read counts of protein coding genes. Color legend. Apparent normal samples: Yellow, DCIS samples: Purple, Paired normal samples: Green, IDC samples: Red **(F)** Venn diagram depicting comparison of differential expression analysis group IDC *vs.* paired normal, IDC *vs.* apparent normal and DCIS *vs.* apparent normal samples **(G)** Venn diagram depicting comparison of differential expression analysis group IDC *vs.* paired normal and IDC *vs.* apparent normal samples **(H)** Venn diagram depicting comparison of differential expression analysis group DCIS *vs.* apparent normal and IDC *vs.* paired normal samples **(I)** Venn diagram depicting comparison of differential expression analysis group IDC *vs.* apparent normal and DCIS *vs.* apparent normal samples

**Supplementary Figure 2. Summary of lncRNA expression profile in IDC *vs.* DCIS. (A)** Volcano plot representing expression pattern in IDC *vs.* DCIS **(B)** Heatmap depicting expression trend of differentially expressed gene in IDC *vs.* DCIS

**Supplementary Figure 3. LncRNA expression profile in various molecular subtype of breast cancer obtained from TCGA dataset using TANRIC platform (A)**RP11-161M6.2 **(B)**ADAMTS9-AS2 **(C)**CTB-92J24.3 **(D)**CTB-131K11.1 **(E)**EPB41L4A-AS1**(F)**FAM83H-AS1 **(G)**LINC01614 **(H)**MIAT **(I)**RP11-295M3.4 **(J)**RP11-490M8.1 **(K)**ST8SIA6-AS1 **(L)**WDFY3-AS2

**Supplementary Figure 4. Expression levels of deregulated lncRNAs in various combination of receptors (ER, PR, HER2) positivity in TCGA dataset (A)** ER+ or PR+ or Her2+ **(B)**ER+ or PR+ along with Her2-**(C)**ER+ and PR-and Her2+ **(D)**ER-or PR-along with Her2+ **(E)**Molecular subtype stratification of validation cohort; Red background: Upregulated lncRNAs and Blue background: Downregulated lncRNAs.

**Supplementary Figure 5. High confidence interaction network (score: 0.7) representing differentially expressed mRNA that are known to co-express with each other as per String analysis and with lncRNA with Pearson correlation coefficient** ≥ **0.9 (A)** RP11-142C4.6 **(B)** RAMP2-AS2 **(C)** RP11-701H24.4 **(D)** PSMB8-AS1

**Supplementary Figure 6. High confidence interaction network (score: 0.7) representing differentially expressed mRNA that are known to co-express with each other as per String analysis and with lncRNA with Pearson correlation coefficient** ≥ **0.9with TINCR**

**Supplementary Figure 7. High confidence interaction network (score: 0.7) representing differentially expressed mRNA that are known to co-express with each other as per String analysis and with lncRNA with Pearson correlation coefficient** ≤ **-0.9with LINC01359**

## Abbreviations

DCIS: ductal carcinoma *in situ*
IDC: invasive ductal carcinoma
NGS: Next generation sequencing
lncRNA: long non-coding RNAs
lincRNA: long intergenic non-coding RNA
TNBC: Triple negative breast cancers
BH: Bonferroni and Benjamini–Hochberg
PCA: Principal component analysis
PCC: Pearson’s correlation coefficient
PCG: Protein coding genes

