## Supplementary figures and images for "Identification of lncRNAs associated with early stage breast cancer and their prognostic implications"

### Supplementary Figure 1

#### Supplementary Figure 1

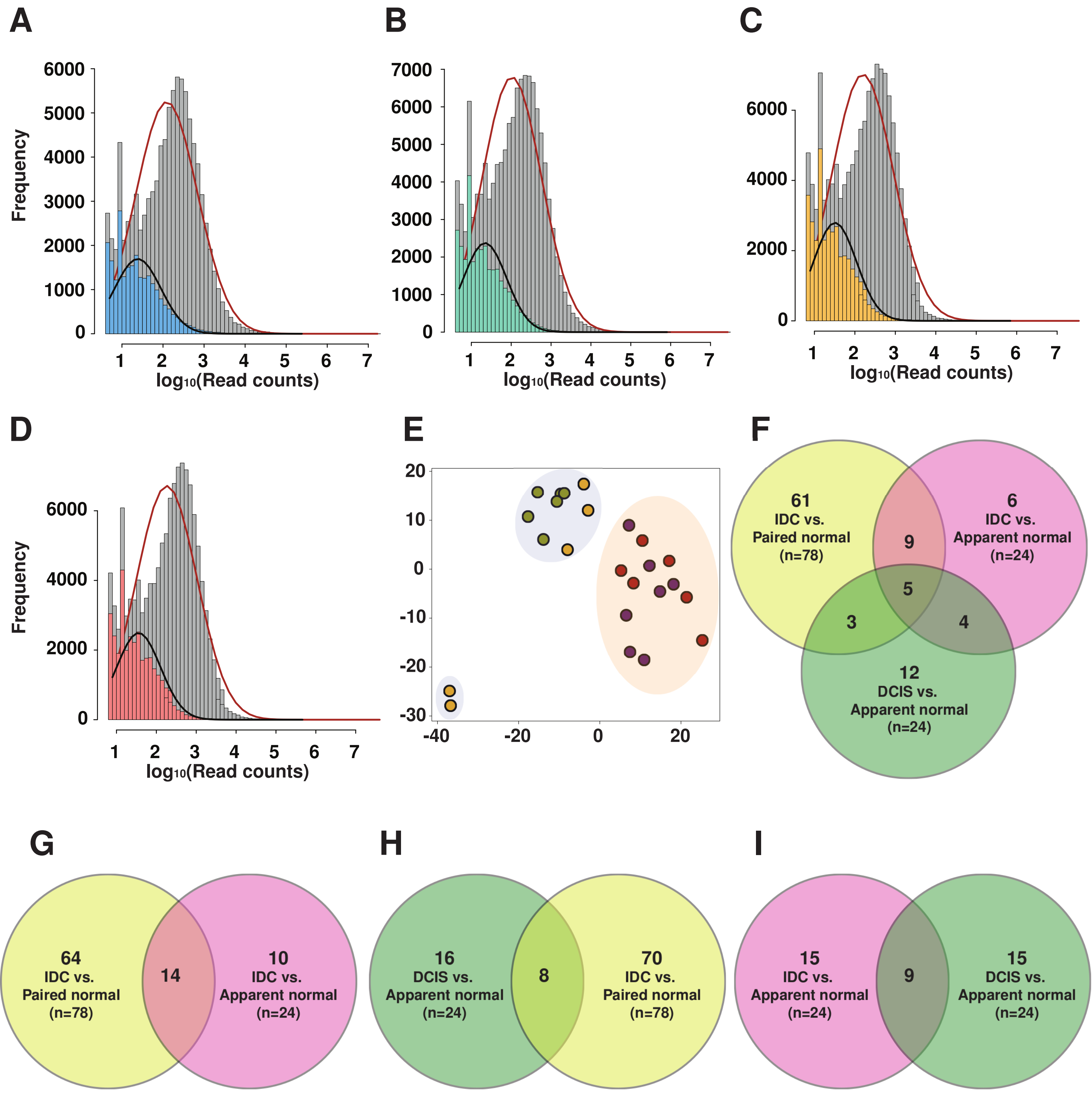

### Supplementary Figure 2

#### Supplementary Figure 2

A

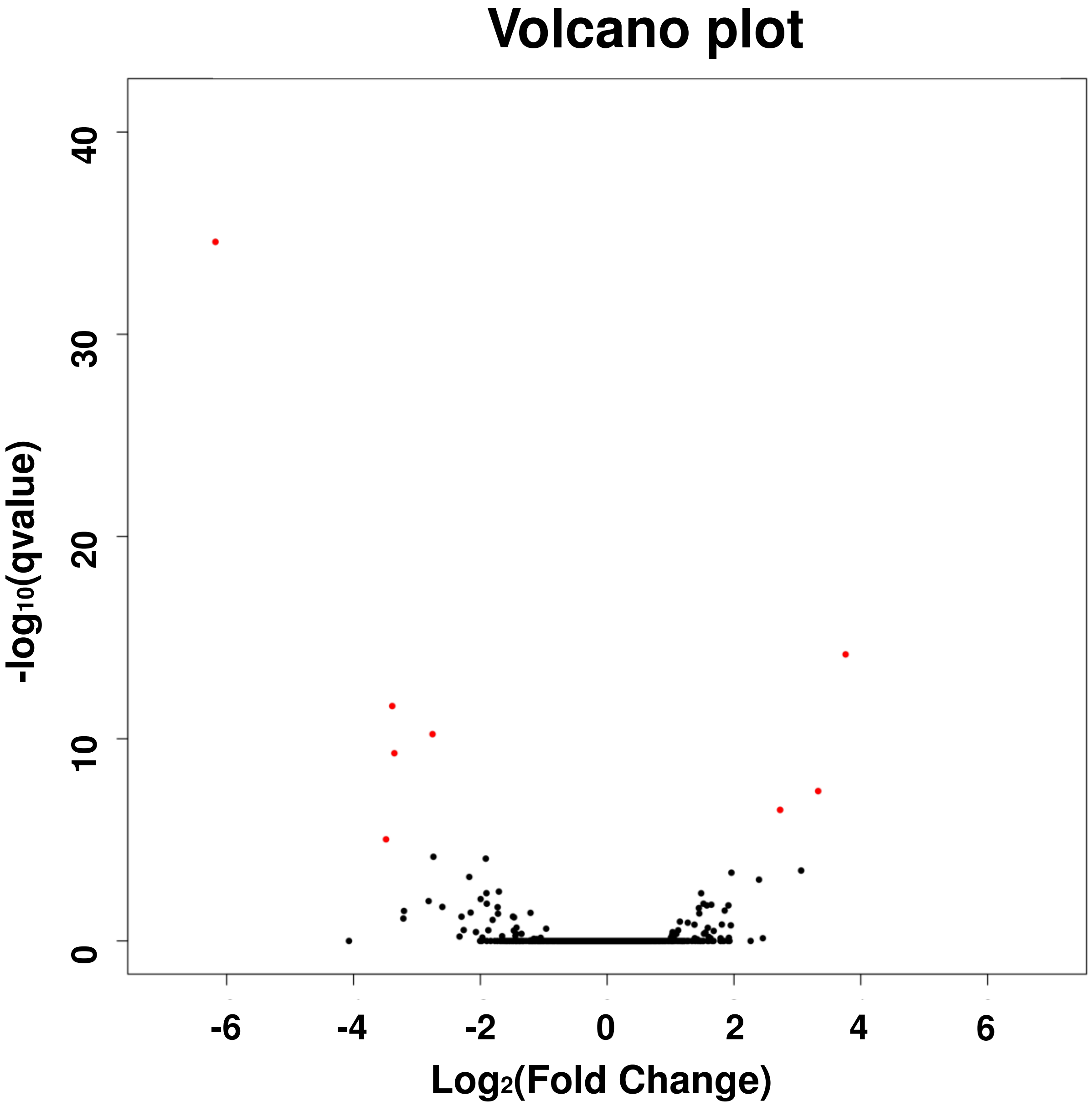


Differentially expressed lncRNA: 12
DE lncRNA exclusive to IDC vs. DCIS: 1

B

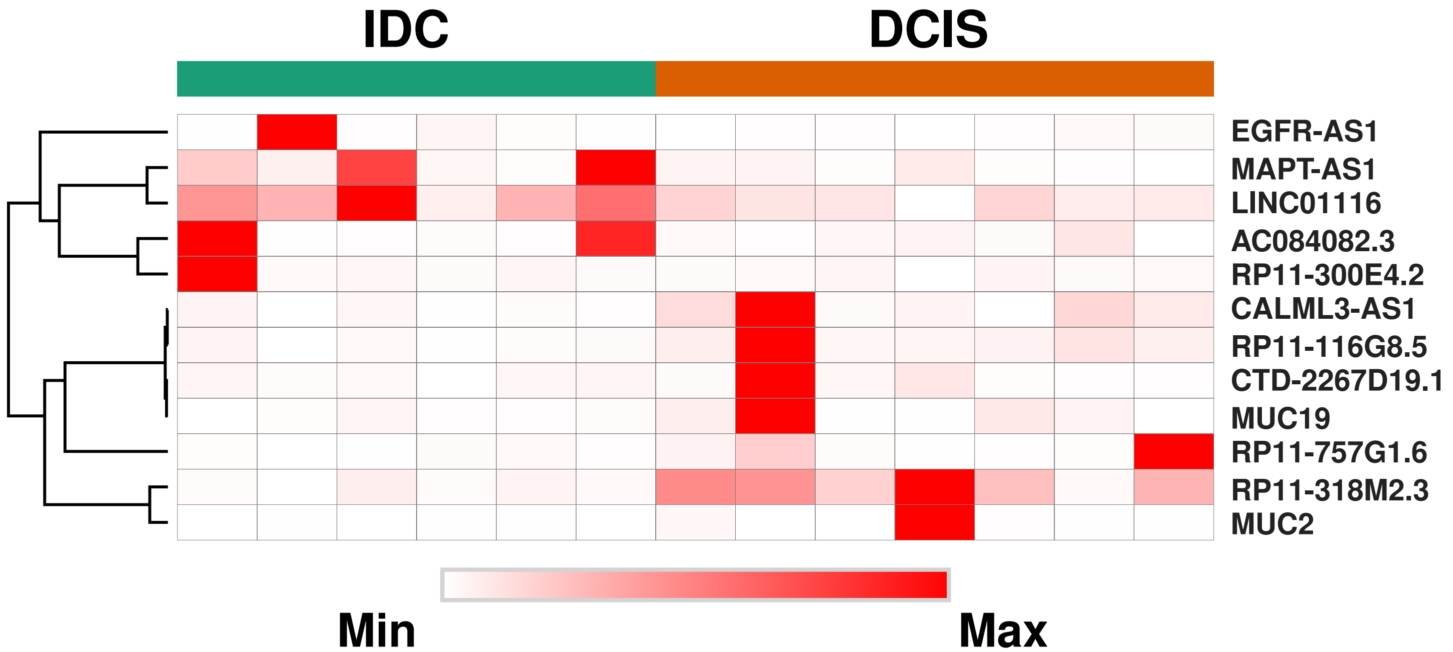

### Supplementary Figure 3

#### Supplementary Figure 3

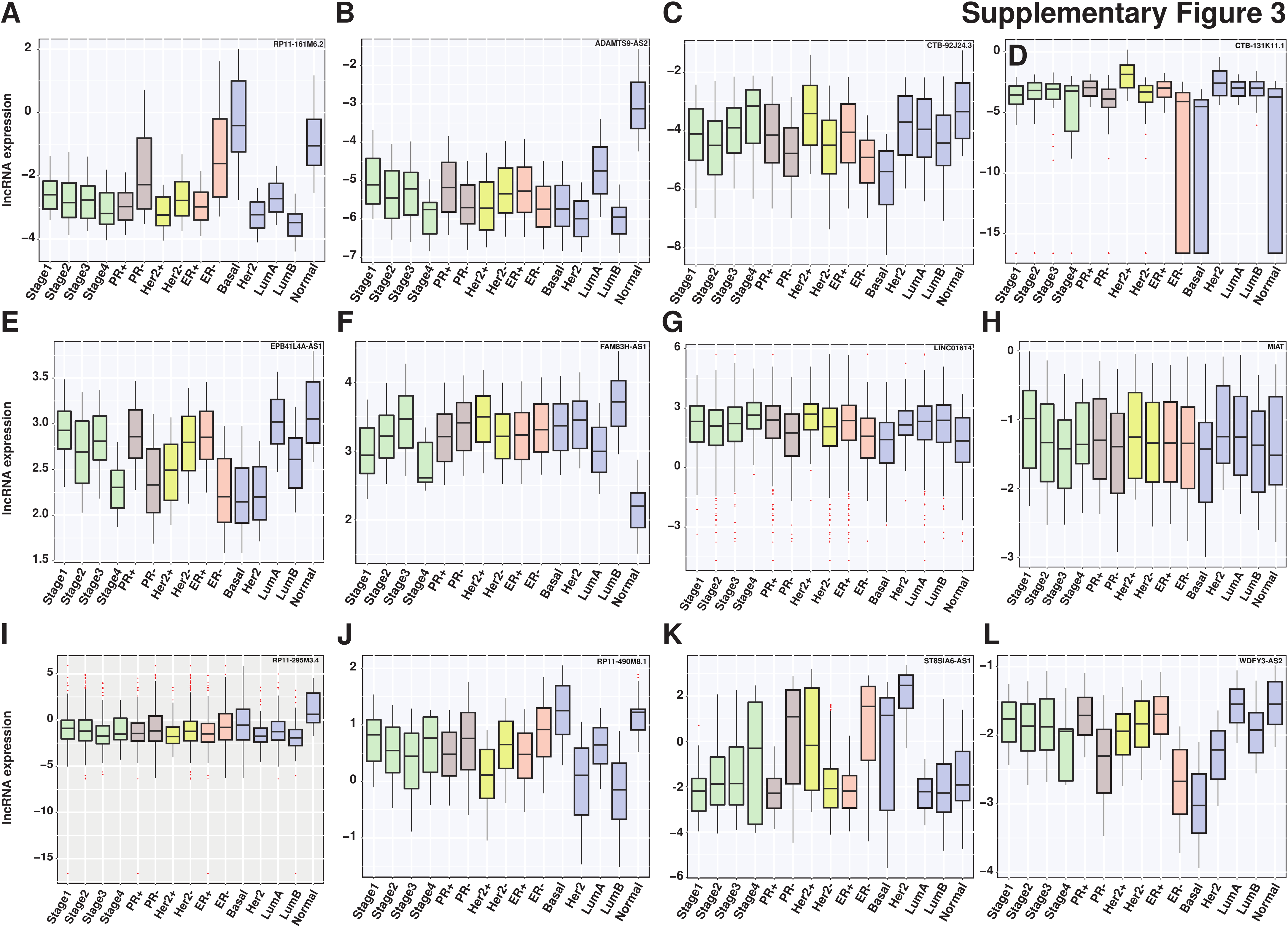

### Supplementary Figure 4

##### Supplementary Figure 4

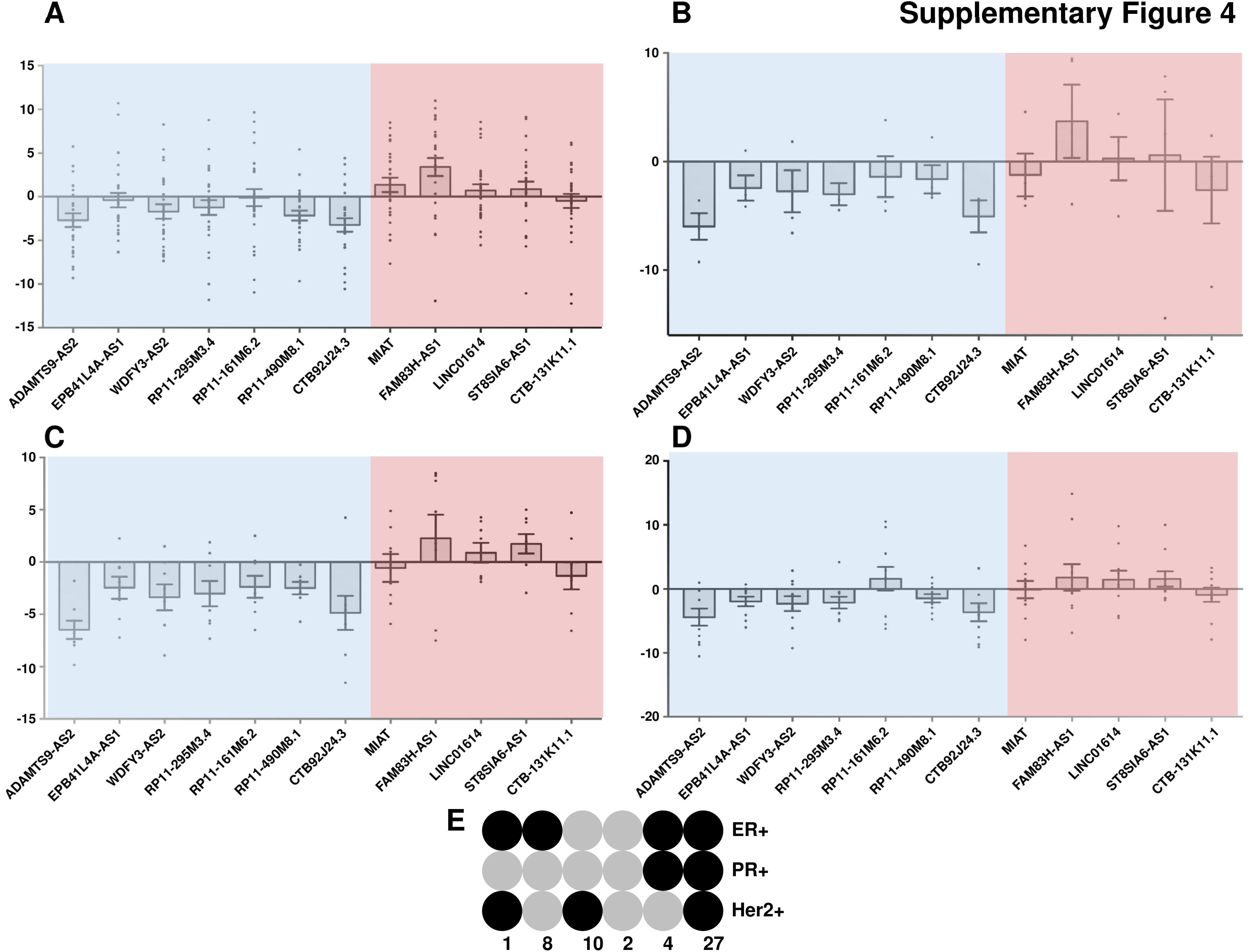

### Supplementary Figure 5

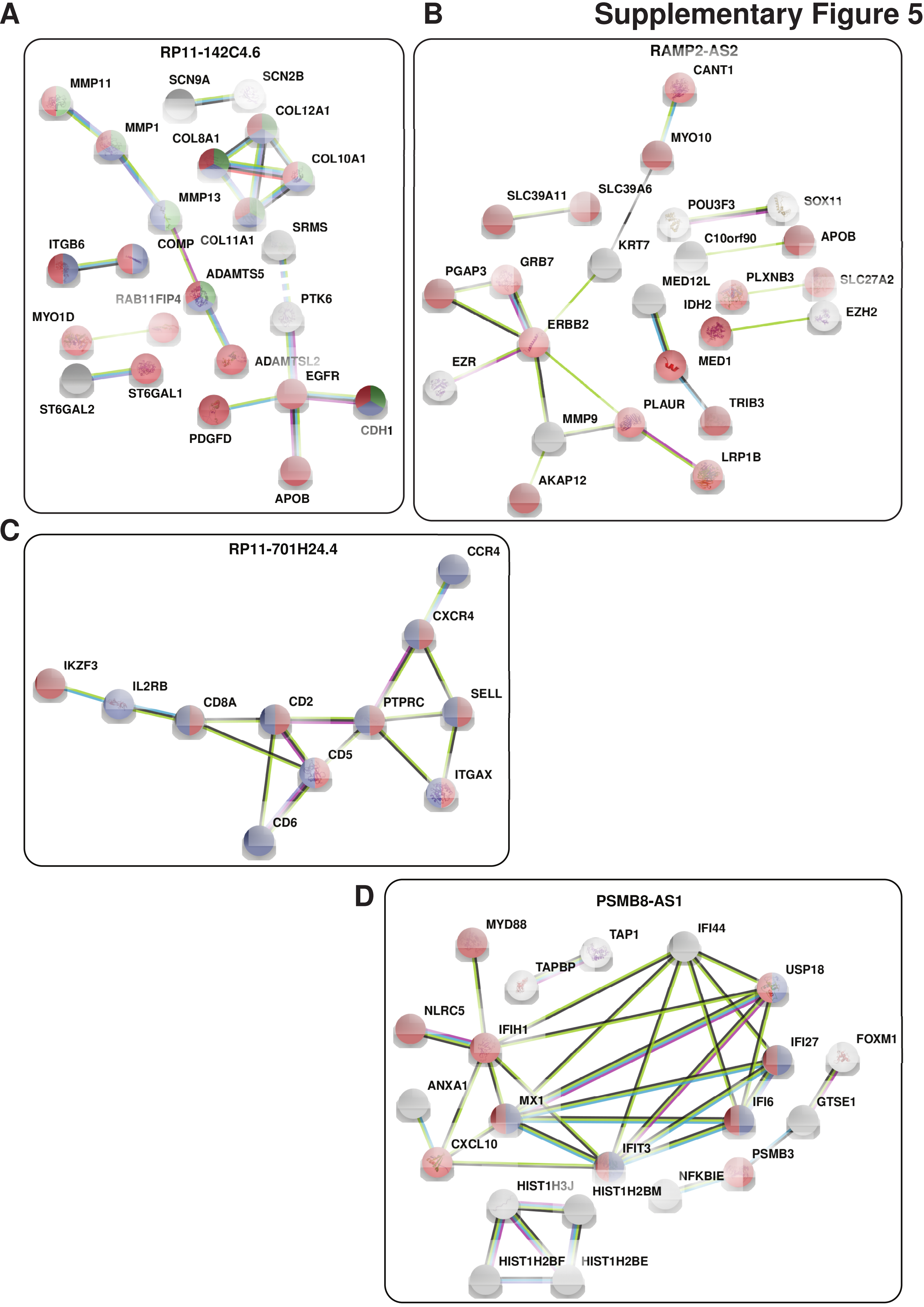

### Supplementary Figure 6

#### Supplementary Figure 6

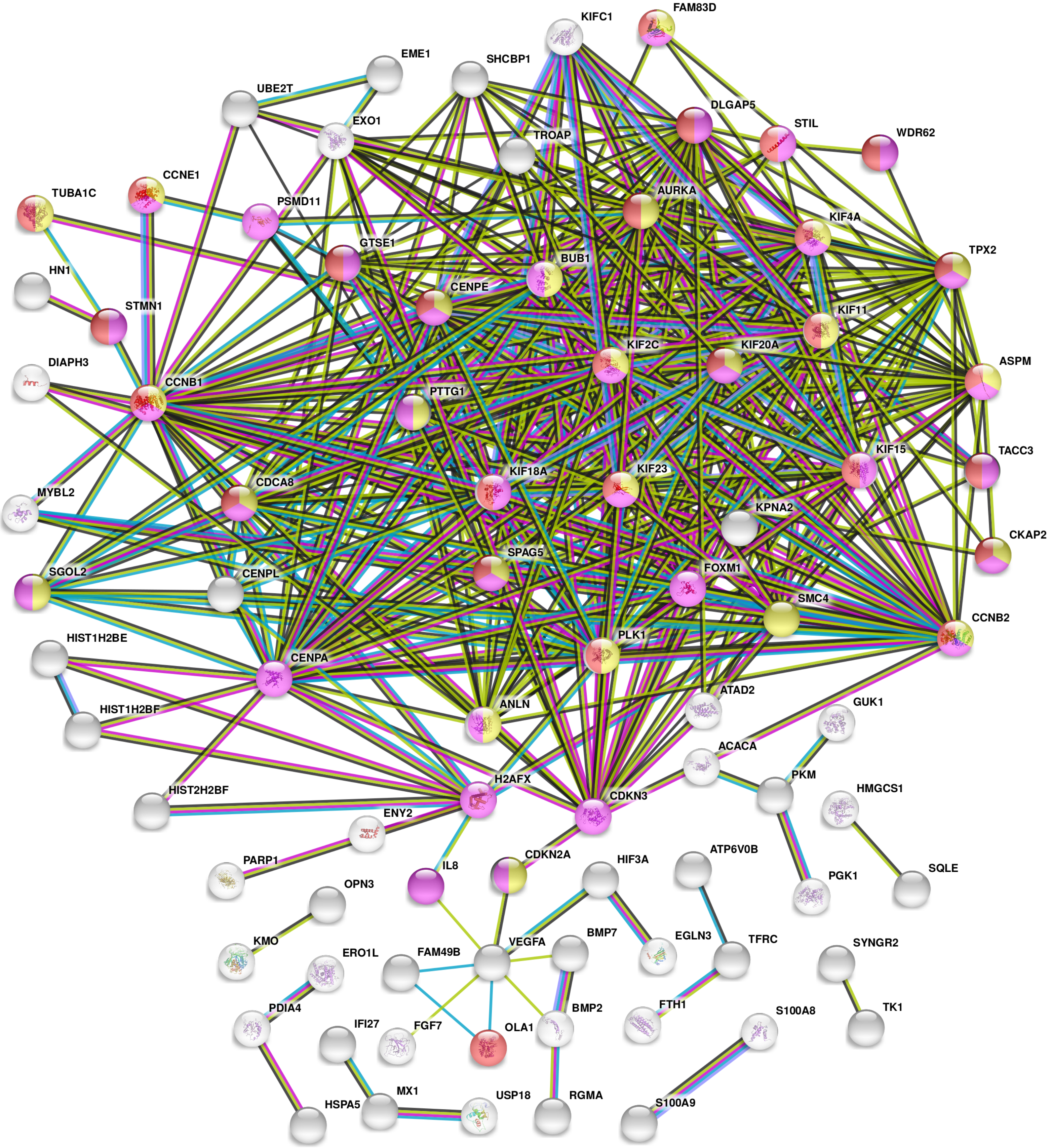

### Supplementary Figure 7

#### Supplementary Figure 7

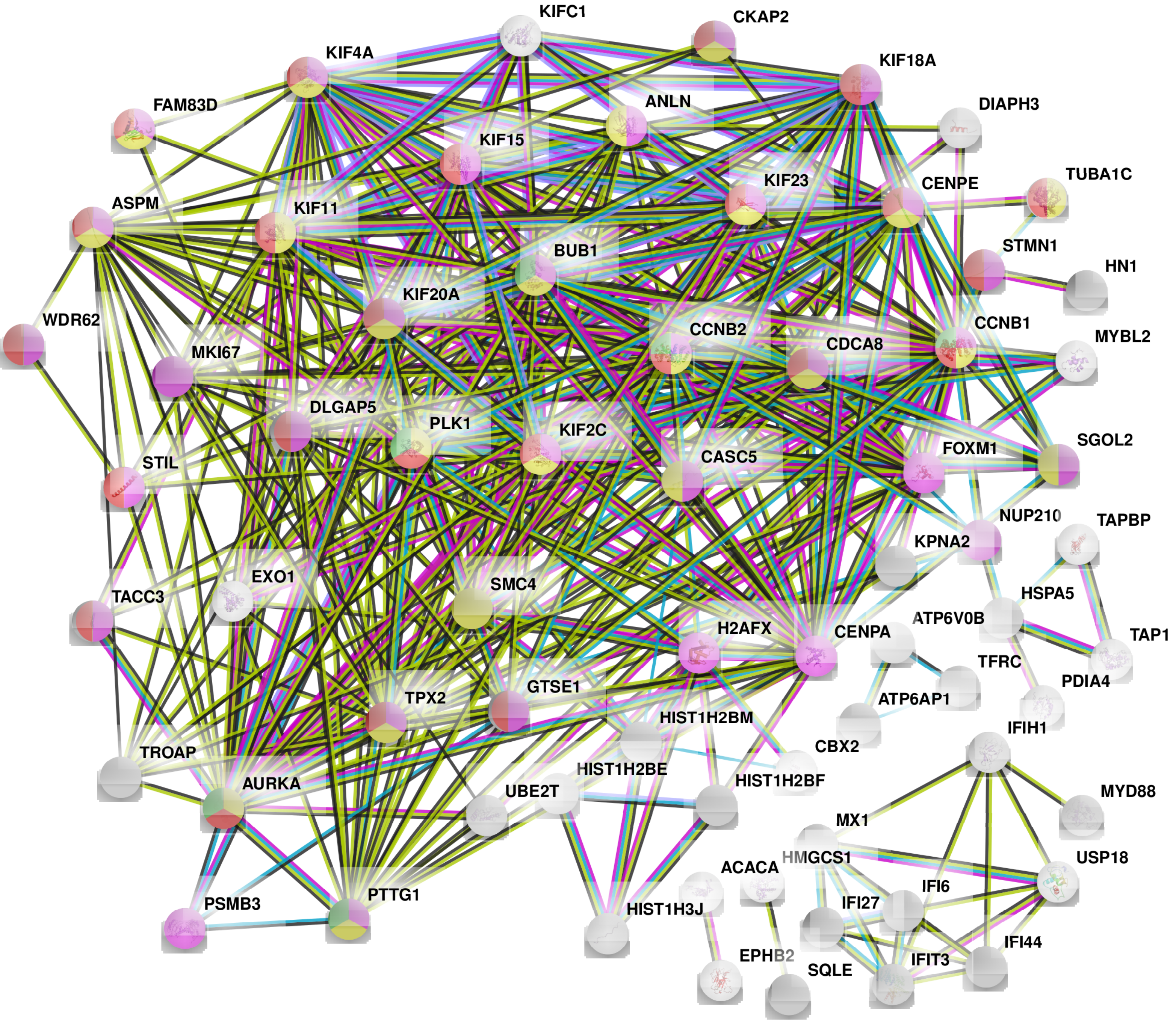
